# Comparison of linear spatial filters for identifying oscillatory activity in multichannel data

**DOI:** 10.1101/097402

**Authors:** Michael X Cohen

## Abstract

**Background:** Large-scale synchronous neural activity produces electrical fields that can be measured by electrodes outside the head, and volume conduction ensures that neural sources can be measured by many electrodes. However, most data analyses in M/EEG research are univariate, meaning each electrode is considered as a separate measurement. Several multivariate linear spatial filtering techniques have been introduced to the cognitive electrophysiology literature, but these techniques are not commonly used; comparisons across filters would be beneficial to the field.

**New method:** The purpose of this paper is to evaluate and compare the performance of several linear spatial filtering techniques, with a focus on those that use generalized eigendecomposition to facilitate dimensionality reduction and signal-to-noise ratio maximization.

**Results:** Simulated and empirical data were used to assess the accuracy, signal-to-noise ratio, and interpretability of the spatial filter results. When the simulated signal is powerful, different spatial filters provide convergent results. However, more subtle signals require carefully selected analysis parameters to obtain optimal results.

**Comparison with existing methods:** Linear spatial filters can be powerful data analysis tools in cognitive electrophysiology, and should be applied more often; on the other hand, spatial filters can latch onto artifacts or produce uninterpretable results.

**Conclusions:** Hypothesis-driven analyses, careful data inspection, and appropriate parameter selection are necessary to obtain high-quality results when using spatial filters.

---

The brain is a complex information-processing organ that produces uncountable dynamics spanning many orders of magnitude in space and in time. A fraction of these dynamics can be measured using electrophysiological techniques such as electroencephalography, magnetoencephalography, and local field potential recordings (EEG, MEG, LFP). These techniques work because large electromagnetic fields spread via volume conduction from brain tissue to electrodes or sensors placed inside or outside the head. The time-varying voltage fluctuations measured by M/EEG and LFP reflect the meso- and macroscopic organization of brain function, and have been linked to myriad sensory, motor, cognitive, and emotional processes.

The vast majority of analysis techniques that are applied in cognitive electrophysiology involve univariate analyses, in the sense that at each step of the analysis, data from only a single electrode are used, and comparisons across electrodes are made qualitatively, e.g., by inspecting topographical distributions. Such methods include short-time Fourier transform, wavelet convolution, filter-Hilbert, multitaper method, empirical mode decomposition, and autoregressive modeling (Cohen, 2014).

But although electrodes are separate, they are not independent. Volume conduction ensures that multiple sources of brain activity contribute to the signal recorded at each electrode. This source-level mixing means that a single electrode is insufficient to isolate different neural generators. Fortunately, different sources have different spatial-spectral-temporal characteristics, and, importantly, the sources are mixed in a linear fashion (Nunez and Srinivasan, 2006). This means that linear spatial filtering techniques can be highly successful at isolating different statistical or anatomical sources. The idea of a spatial filter is to use weighted combinations of electrode activity to identify patterns in the data that are difficult to observe in the unfiltered data. The weights are defined by statistical and/or anatomical criteria, and are guided by the goal of isolating sources of variance in multichannel data, identifying putative anatomical generators, or denoising data during preprocessing. Commonly used classes of spatial filtering include independent components analysis (ICA), the surface Laplacian, and source localization methods such as dipole-fitting, LORETA, and beamforming, although there are many other spatial filtering methods.

Because the number of electrodes used in electrophysiology is increasing (Stevenson and Kording, 2011), and because there is growing awareness that the spatial dimension contains a significant amount of information that can be used for denoising (e.g., removing noise components in ICA) or for analysis (e.g., increasing signal-to-noise ratio or applying various machine-learning algorithms to “decode” brain states) (King et al., 2014; Onton et al., 2006; Vigario and Oja, 2008), spatial filtering techniques are likely to become increasingly important in cognitive electrophysiology.

On the other hand, linear spatial filters are not widely used in cognitive electrophysiology research. There are papers that detail the mathematical backbone of linear spatial filtering (Congedo et al., 2008; Parra et al., 2005), and other papers that offer motivations and case-studies for applying such analyses (Cunningham and Yu, 2014; Pang et al., 2016; Vigario and Oja, 2008). But these papers are often of limited practical use to researchers seeking nontechnical explanations, comparisons across filters, or illustrations of use in real data. Linear spatial filters are applied more widely in the brain-computer-interface field, and several papers have provided rigorous comparisons amongst different linear spatial filtering techniques (Foldes and Taylor, 2011; Jochumsen et al., 2015). These studies highlight the usefulness of spatial filtering and component extraction procedures, but their direct relevance to cognitive electrophysiology is less clear. In particular, cognitive electrophysiology studies often involve offline data analyses that aim to identify neural activity patterns that are frequency-band-limited, temporally brief, and correlated with experiment conditions and individual differences.

Therefore, the purpose of this study is to compare six linear spatial filters on simulated and on real data, with the focus on using dimensionality-reduction to identify narrow-frequency-band components in multichannel recordings. The goals of the spatial filters were to maximize frequency-specific signal-to-noise ratio, reconstruct the time course of simulated activity, without overfitting noise. There are too many spatial filters to provide a complete comparison in one paper; the methods discussed here are (1) based on a mathematical procedure called generalized eigendecomposition (explained below) and (2) are amenable to dimensionality-reduction-based analyses, in which the goal is to obtain a single component (or perhaps a small number of components) that is analyzed instead of individual electrodes. This latter motivation can be contrasted with other spatial filters that result in the same dimensionality (e.g., the surface Laplacian) or in increased dimensionality (e.g., distributed source localization). The following methods were compared (see Methods for explanations): principal components analysis, spatial-spectral decomposition, generalized eigendecomposition-broadband, joint decorrelation, best-electrode, and independent components analysis. The latter two are actually not based on generalized eigendecomposition, but are included here as reference because they are commonly used in cognitive electrophysiology.

## Methods and materials

### Overview of spatial filtering methods

There are many excellent sources that discuss the mathematics underlying linear spatial filtering techniques; interested readers may start with these (Congedo et al., 2008; de Cheveigné and Parra, 2014; Parra et al., 2005; Sarela and Valpola, 2005). Only descriptive overviews are provided here (see also Figure 1) for conceptual understanding.

**Figure 1.**
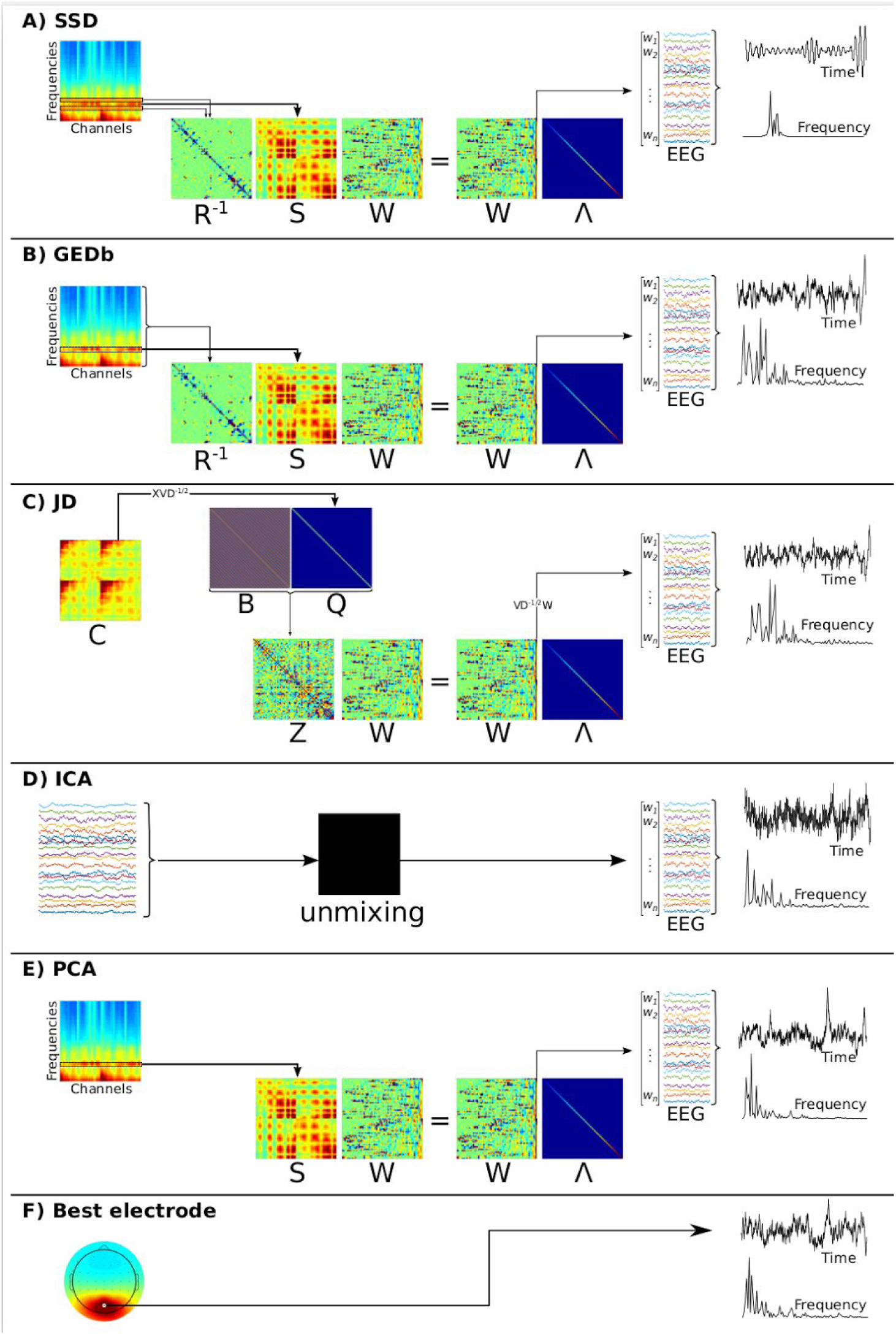
Graphical overview of the linear spatial filters investigated here. All matrices are size channel-by-channel unless marked otherwise. Matrix **S** (“signal”) is a covariance matrix from data bandpass filtered at the target frequency; matrix **R^-1^** (“reference”) is the inverse of a covariance matrix from data bandpass filtered at a flanking frequency, or from broadband data; matrix **W** contains eigenvectors and matrix ***Λ*** contains their corresponding eigenvalues on the diagonal. The myriad steps of ICA (panel D) are beyond the scope of visual illustration here; interested readers are referred to Figure 2 of Rutledge and Bouveresse (2013).

Four of the spatial filters considered here rely on generalized eigendecomposition. When matrix **A** is a covariance matrix, its eigendecomposition **AW=WΛ** produces a set of vectors in **W** that point along the principal axes of covariance in the data from which **A** is defined, and the diagonal matrix **Λ** indicates the importance of each direction. A generalized eigendecomposition of two covariance matrices **A** and **B** can be used to find a set of vectors that characterize the distances between those matrices, and is written as **AW=BWΛ**. One can think of rewriting this equation as **B^-1^AW=WΛ**, in other words, find the vectors W that maximize the “ratio” of **A** to **B**. When covariance matrix **A** is computed from to-be-enhanced activity and covariance matrix **B** is computed from to-be-suppressed activity, the vectors in matrix **W** are the spatial filters that best separate **A** from **B**, and the diagonal elements of matrix **Λ** indicate the importance of those vectors. Unlike with principal components analysis, the columns of matrix **W** are generally not orthogonal, because the matrix **B^-1^A** is generally not symmetric.

Temporal bandpass filtering of EEG data was done via convolution with a frequency-domain Gaussian kernel. The full-width at half-maximum (FWHM) was 2 Hz. When spatial filters were applied to broadband data, the component time series were filtered around the peak frequency at 5 Hz FWHM for comparison with the simulated signal. These filter parameters are important predictors of the quality of the spatial filter; the effects of parameter selections on the components are systematically evaluated in the Results section.

#### Spatiospectral decomposition (SSD)

SSD (Nikulin et al., 2011) involves computing a generalized eigendecomposition of two covariance matrices, one corresponding to “signal” and one corresponding to “noise” (or “reference”). The eigenvector with the largest corresponding eigenvalue is taken as a spatial filter that maximally differentiates the two covariance matrices (Figure 1a). In SSD, the signal covariance matrix is computed from bandpass filtered data at the frequency of interest, while the noise covariance matrix is computed from bandpass filtered data at neighboring frequencies. The spatial filter is then applied to the bandpass filtered data used to construct the “signal” covariance. Bandpass filtering was done via 2nd order Butterworth filters instead of convolution with a Gaussian kernel. The Butterworth filter was unable to resolve low frequencies, which is why the SSD data lines do not go down to 2 Hz in the figures. MATLAB code was used from the original publication (https://github.com/svendaehne/matlab_SPoC/tree/master/SSD).

#### Generalized eigendecomposition, broadband (GEDb)

The procedure is similar to SSD except (1) the “reference” covariance matrix is generated from broadband data instead of from neighboring frequencies, (2) the spatial filter is applied to the broadband data instead of to the bandpass filtered data, and (3) the temporal bandpass filter is implemented as convolution with a frequency-domain Gaussian, which allows greater frequency specificity and reduced risk of poor filter design. See Figure 1b and de (Cheveigné et al., 2015) for more details.

#### Bias-filter joint decorrelation (JD)

See (de Cheveigné and Parra, 2014) for details of joint decorrelation; following is a brief description. (1) The data are “sphered” meaning that variance along each dimension is normalized. This is achieved by reconstructing the data as **Y=XVD^-1/2^**, where **X** is the time-by-channels data matrix, **V** is a matrix of eigenvectors, **D** is a matrix with eigenvalues on the diagonal, and ^-1/2^ indices the matrix square root of the inverse. The purpose of sphering the data is to remove the intrinsic variance structure (Figure 1c depicts **Y**’s covariance matrix **Q**, which illustrates that sphered data have a diagonal covariance matrix). (2) A “bias filter” is created that left-multiplies the sphered data (**BY^T^**) (Figure 1c illustrates **Q** instead of **Y**, because **Y** is too large to fit in the figure). Here, the bias filter was a Toeplitz matrix constructed from a sine wave. (3) An eigendecomposition is computed on the covariance matrix (matrix **Z**) of the sphered and filtered data, giving eigenvector and eigenvalue matrices **W** and **Λ**. (4) Finally, the spatial filters are defined as **VD^-1/2^W**. The importance of each vector is defined by the relative sizes of the eigenvalues from the eigendecomposition of the bias-filter-convolved data (also called the variance ratio). Importantly, the spatial filter is then applied to the raw data, not to the sphered data. The topographical maps are taken from the inverse transpose of the filter matrix (Haufe et al. 2014).

#### Best-electrode

One could think of a single electrode loosely as a spatial filter where the weights are zero for N-1 electrodes and one for the selected electrode. In this study, it served as a “straw-man” reference: Any useful spatial filter should outperform the best-electrode approach, if for no other reason than overfitting data. The analyzed electrode (termed “best electrode”) was selected based on the electrode that had maximal power at the frequency of the simulated time series.

#### Principal components analysis (PCA)

PCA involves an eigendecomposition of a covariance matrix, and the M-by-1 (M is the number of channels) eigenvector with the largest eigenvalue is taken as the spatial filter. Temporal bandpass filtering provides frequency specificity of the covariance matrix. PCA is also a straw-man reference: pairwise orthogonal eigenvectors are generally suboptimal in neuroscience data, because the large-scale dynamics of the brain recorded by M/EEG are not orthogonal. Furthermore, with PCA one specifies only the characteristic of the data to be maximized (here, the covariance of narrowband-filtered data); there is no "reference” characteristic to provide a minimization objective.

#### Independent components analysis (ICA)

ICA has the goal of unmixing (rather than decorrelating as with PCA) multichannel datasets (Rutledge and Jouan-Rimbaud, 2013). ICA is widely used in neuroscience, most often as a tool to facilitate data cleaning by identifying and removing components that capture non-brain dynamics such as oculomotor, muscle, or heart beats. There are several ICA algorithms, and although different methods have different implementations and assumptions, they can produce similar decompositions of EEG data (Delorme et al., 2012). The Jade algorithm (Cardoso, 1999) was applied here because it is fast and is deterministic (that is, repeated calls on the same data will produce the same components). Briefly, the Jade algorithm is based on the assumption that pure signals tend to have non-Gaussian distributions while noise or random mixtures of signals tend to have Gaussian distributions; the goal is then to determine weighted combinations of electrodes that maximize non-Gaussian distributions. The Jade algorithm was applied to bandpass filtered data, and the 30 largest components were extracted. The component with the highest power at the simulated frequency (this was obtained by computing an FFT of the IC time series and extracting power at the frequency of the simulated signal for all components) was selected for subsequent comparisons.

### Simulated and empirical data

A forward model was created using the brainstorm toolbox in Matlab (Tadel et al., 2011), comprising 2004 dipoles distributed throughout the cortex. Each dipole was given a time series of smoothed random numbers. Two dipoles were selected because they projected to central posterior electrodes, and their time courses were simulated as sine waves with time-varying amplitudes and frequencies. The “target” oscillation was simulated at frequencies between 2 and 80 Hz, with one frequency present per simulation run. An additional (“distractor”) frequency was added to the second dipole at a frequency between 1 and 6 Hz higher than the target frequency (randomly selected on each simulation run). Thirty seconds of continuous data were simulated at a sampling rate of 1024 Hz. The first and last 500 ms were excluded from data analyses.

The goal of the spatial filters was to recover the time series of the target dipole. After simulating the time series, data from all dipoles were then projected to the scalp. In different sets of simulations, the projected dipole data were either summed with random noise, or were added to real EEG data that was measured from a human volunteer during an eyes-open resting condition. EEG data were recorded from 64 electrodes using Biosemi equipment (www.biosemi.com) with a sampling rate of 1024 Hz. In another set of simulations with the empirical EEG, data from electrode O1 were replaced with random noise. The purpose of this was to determine the sensitivity of different spatial filters to a noisy or broken electrode, which is a frequent annoyance in EEG research.

Task-related empirical data were included to provide a proof-of-principle application in real data that contain real physiological activity and noise sources. Data were taken from a single subject from a previously published study (Cohen, 2015). The subject performed a Flankers task, in which the task was to identify the central letter from flanking letters (e.g., “T T I T T”). Trials containing excessive EEG artifacts were manually rejected, but other artifacts (e.g., oculomotor, EMG, electrode noise) were not removed using independent components analysis. The reasons for not subtracting noise components were (1) to keep the data matrices at full-rank, and (2) to test whether the spatial filters were robust to typical EEG artifacts. The analyzed dataset includes data from 64 electrodes (512 Hz) and 250 trials, one-half of which were correct responses and one-half were errors. Additional experimental details can be found in Cohen, 2015.

Although the “true” underlying features of empirical data are unknown, strong a priori expectations can be made on the basis of many published studies (Cavanagh et al., 2009; Luu et al., 2004; van Driel et al., 2012). Averaging both conditions relative to baseline should reveal an increase in delta-theta power at midfrontal regions, and a decrease in alpha-beta power in motor and visual/parietal regions; and the contrast of error vs. correct should yield an increase in midfrontal delta-theta power.

### Evaluating spatial filter performance

The spatial filters were evaluated using two criteria. First was the correlation between the spatially filtered result and the simulated dipole signal. R^2^ was used instead of the raw correlation coefficient because the sign was occasionally flipped. Sign-flipping occurs often in eigendecomposition-based methods, because the direction of an eigenvector is not necessarily meaningful. For visual comparison, the signs of the topographical maps were adjusted so that the largest-magnitude electrode had a positive sign.

The second evaluation was spectral signal-to-noise ratio (SNR). The power spectrum of the component time series was extracted via the FFT. SNR was defined as the ratio of power at the peak frequency to the average power of the surrounding ±5 Hz, but excluding ±1 Hz around the peak (Cohen and Gulbinaite, 2016). Because the SNR peak was defined by the frequency of the simulation, SNRs>1 can be expected; the important comparison is of SNRs across different filters.

Simulations were repeated 20 times using different random seeds. For many comparisons, statistics were not applied because, e.g., all 20 results from one filter were much larger than all 20 results from another filters. When significant differences were not visually obvious, ANOVAs were applied in Matlab, with “filter type” and “noise condition” (O1 comprising or not comprising noise) being the independent variable factors.

The sensitivity of the spatial filters to their parameters (primarily the widths of the temporal bandpass filters) was also investigated by systematically varying filter parameters and the number of simulated electrodes.

MATLAB scripts and data can be downloaded from mikexcohen.com/data.

## Results

### Recovering simulated oscillations in noise

In the first set of simulations, lowpass filtered random noise was generated at 2004 dipoles in the cortex. Data from two dipoles were replaced with a signal comprising a sine wave with temporal nonstationarities in amplitude and frequency. The activities from all dipoles were then projected onto 64 scalp EEG channel locations, and analyses were conducted on the simulated EEG time series.

Figure 2a shows results for fits of the component time courses to the time courses of the target simulated dipole. The three methods based on generalized eigendecomposition performed well (R^2^>.8), with R^2^ values decreasing from GEDb to JD to SSD (1-way ANOVA F_2,57_=1615; all pairwise t-tests significant at p<.001). The generalized eigendecomposition-based methods and ICA were fairly robust to replacing data from electrode O1 with pure noise. Best-electrode and PCA suffered more from the noisy electrode; on closer inspection, the poor performance of the noisy electrode was attributable to the automatic electrode selection procedure, which was based on maximizing power at the target frequency. The broadband noise power at O1 was often higher than the target frequency power at POz. The best-electrode and PCA results are nearly perfectly overlapping in Figure 2a2 and 2b2, because the first principal component was almost exclusively driven by electrode O1.

**Figure 2.**
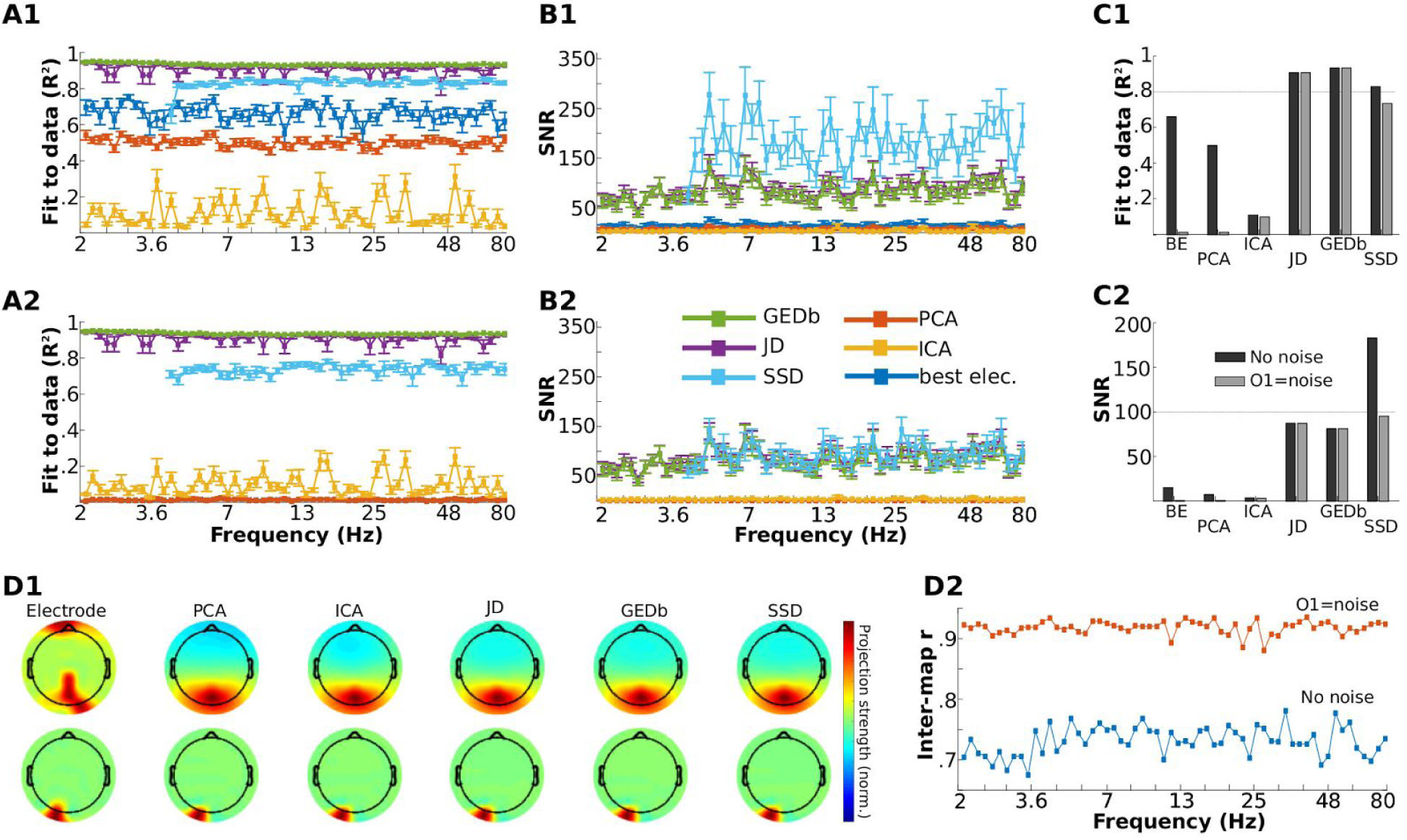
Results of different spatial filters applied to noise data (“EEG”) in order to identify and reconstruct an embedded signal. All results show mean and standard error over 20 simulations with different random noise seeds. Analyses were done twice on each simulation, once projecting noise from dipoles to EEG electrode locations, and another using the same data but replacing electrode O1 (close to the maximal projection of the source) with random noise. Panel A shows the R^2^ fit of the filtered component to the simulated time series (A1 is for the unadulterated simulations, A2 is for the O1=noise simulations). Panel B shows spectral SNR at the simulated frequency. Panel C shows the results from panels A and B averaged over frequencies. Horizontal lines are drawn at R^2^=.8 and SNR=100. Panel D1 shows the topographical maps of the data and various filter projections (“Electrode” refers to frequency-specific power at each electrode). Topographical maps here and in subsequent figures were normalized to facilitate visual comparison; relative color values can be compared across spatial filters. All frequencies had similar topographical maps. Panel D2 shows the average inter-filter topographical correlations.

SNR showed comparable performance as fits to the simulated time course (Figure 2b), with the three generalized eigendecomposition methods performing favorably, and SSD (without the noise-O1 replacement) showing the highest SNR. JD SNR was slightly but significantly higher than GEDb SNR (t_38_=2.65, p=.015). SSD was more sensitive to replacing O1 activity with noise. Nonetheless, even the seemingly “poor” performance of the best-electrode approach was adequate, with SNRs averaging around 15, and going up to 30 in some cases. Figure 2c provides an overview of the results averaged over target frequencies.

The topographies of power and the spatial filter forward model projections are shown in Figure 2d. Average spatial correlations amongst all filters — which indicates the consistency of identifying dynamics across different filters — were around .75 without noise and around .9 with the noisy electrode. It is interesting to observe that the noisy electrode severely impacted the visual interpretability of the spatial filter topographical projections, but the fit to the simulated signal and the SNRs were often only mildly affected.

This set of simulations was repeated without adding the “distractor” frequency; the results (not shown) were all qualitatively the same as those reported above. In some sense, this is not surprising given the temporal filtering used to create the spatial filters, which was narrow enough to exclude the distractor frequency in many cases. Nonetheless, it demonstrates the utility of applying *spatiotemporal* filtering for component isolation, as opposed to spatial-only filtering.

#### Recovering simulated oscillations in real data

Because simulated noise lack physiological characteristics of real EEG data, the evaluations described above were repeated but adding the simulated signal on top of real EEG data (taken from a resting-state recording) instead of noise. Results are shown in Figure 3.

**Figure 3.**
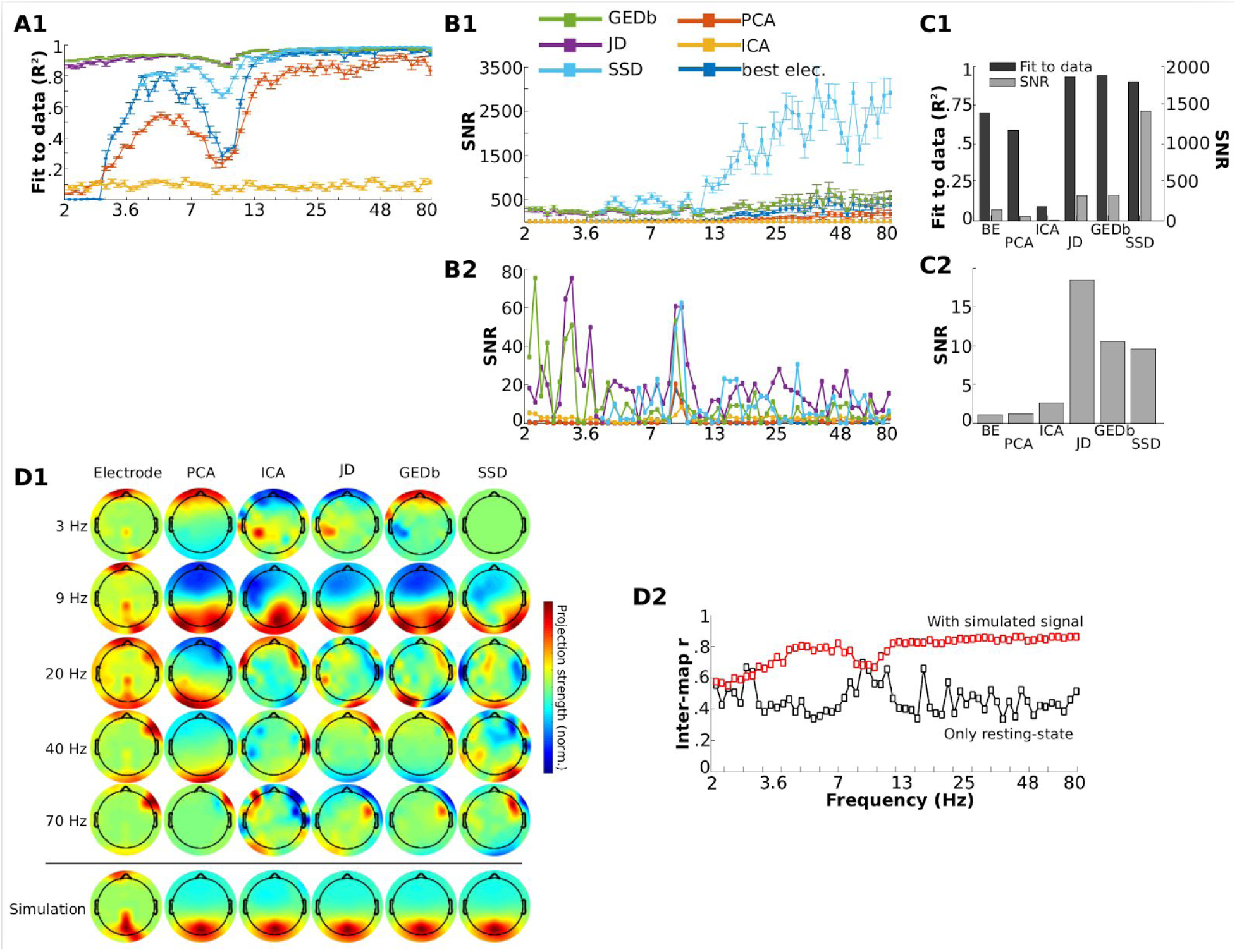
Similar to Figure 2, except that the simulated time course was added to real resting-state EEG data. In panels A, B, and C, subpanel “1” is for the analyses with the simulated signal added, and subpanel “2” is with no simulated signal added (thus reflecting only the empirical resting-state data).

GEDb and JD provided the best fits to the simulated signal, with R^2^’s above .85 for all frequencies. SSD, best-electrode, and PCA provided good fits for higher frequencies. All methods had a relative dip in the alpha-band, attributable to the strong endogenous alpha activity overpowering the simulated signal (simulated signal amplitude was the same for all frequencies and was thus relatively lower where the EEG power was relatively greater). The frequency-averaged R^2^’s decreased from GEDb to JD to SSD (one-way ANOVA F_2,57_=570, all pairwise t-tests significant at p<.001).

SNRs were comparable for the three eigendecomposition-based methods below around 15 Hz. For higher frequencies, SSD SNR continued to increase, leading to a higher frequency-averaged SNR. The frequency-averaged SNR was higher for SSD than for the other methods (one-way ANOVA F_2,57_=595, JD and GEDb were not significant different from each other, p=.57).

Spatial filters were next applied to the data without adding the simulated sine wave (thus, only real EEG data). All methods produced an SNR peak in the alpha band, as well as some additional lower frequency peaks. However, different spatial filtering methods identified different topographical patterns over frequencies, leading to an overall spatial correlation across maps of around .5. JD had the highest average SNR.

### Different parameters within spatial filters

The purpose of this set of simulations was to examine the effects of temporal filter parameters on the performance of the spatial filters. (The amplitude of the simulated signal was decreased by an order of magnitude to avoid ceiling effects in reconstruction accuracy.)

The main parameter for JD is the width of the temporal filter used to reconstruct the component after the spatial filter has been applied to the broadband data. Figure 4a shows that while very narrow filters are suboptimal (because they also attenuate the nonstationarities contained in the simulated signal), a range of wider filters provided reasonably accurate fits. The bias filter could be constructed either as a sine or as a cosine wave. The sine wave filter provided a better fit for these simulations, highlighting that the bias filter in JD can be phase-dependent.

**Figure 4.**
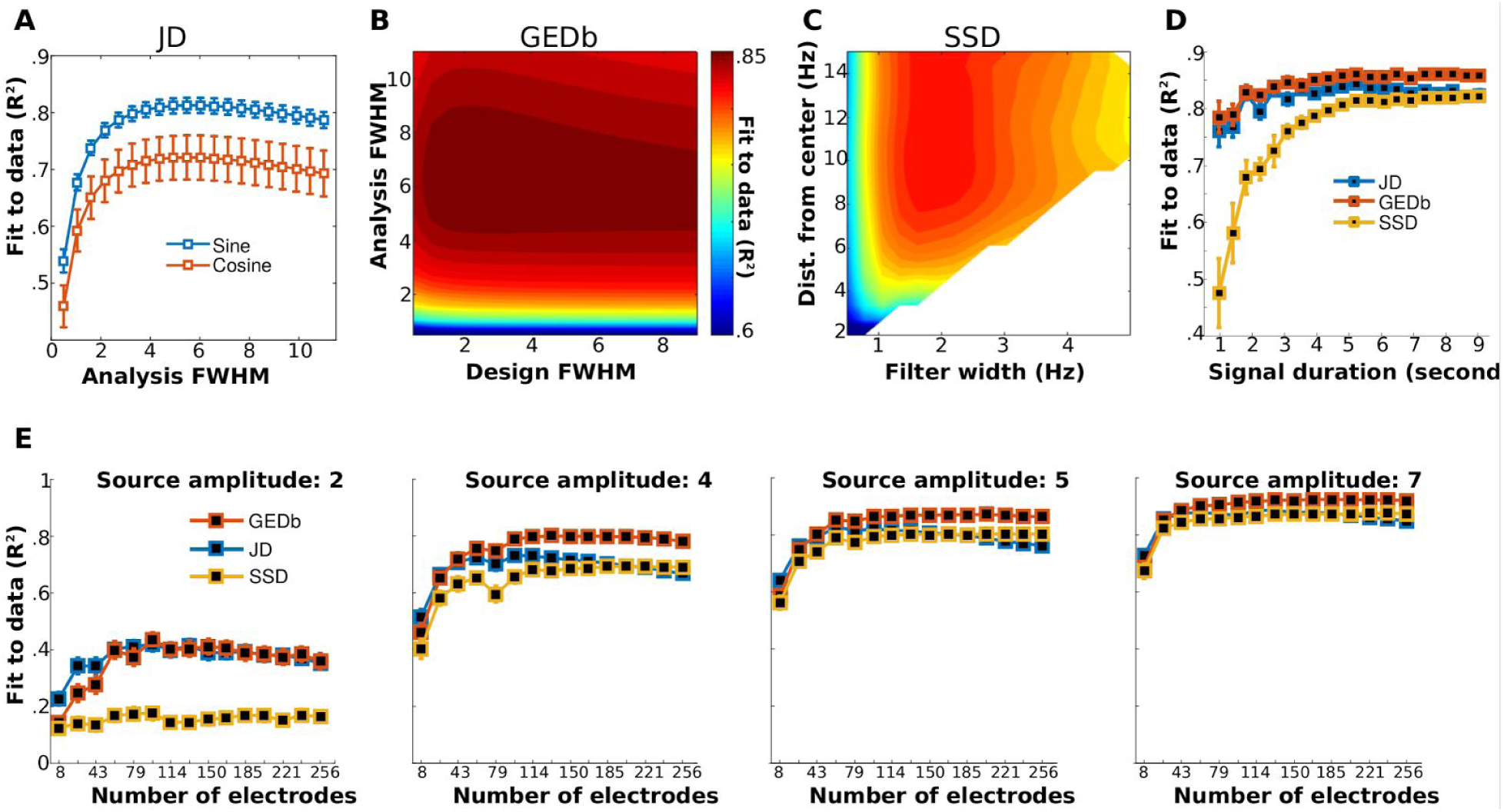
Effects of temporal filter parameters on fits to the simulated data (R^2^) for JD (A), GEDb (B), and SSD (C). Panels B and C have the same color scaling. Results show the average (and standard error of the means for panels A and D) across 20 runs with the same parameters but different random seeds for simulating data. Panel D shows the fit to the simulated data as a function of the duration of the time series. Panel E shows R^2^ fits for different numbers of electrodes. Each panel corresponds to a different dipole signal strength.

GEDb has two temporal filter parameters, the FWHM used when creating the filter (“design FWHM” in Figure 4b), and the FWHM used when bandpass filtering the component time series (“analysis FWHM” in Figure 4b). In the simulations performed here, the optimal parameter-pair was around 3 Hz and 6 Hz for the design and analysis FWHMs. However, fits were generally high for most parameter pairs sampled, except for very narrow analysis FWHM (such stringent filtering attenuated the nonstationarities in the simulated signal).

SSD has two temporal filter parameters, the width of the flanking frequencies and their distance from the target frequency. The full parameter space could not be sampled, because the implementation from Nikulin et al. does not allow the flanking frequencies to overlap with the target frequency. Results show that in this simulation, the best fit was obtained from relatively narrow spectral flankers (~2 Hz width) that are relatively far from the target frequency (>6 Hz, Figure 4c).

The sensitivity to the length of the time window that was used to construct the covariance matrices was also examined. Figure 4D shows that all three tested filters provided strong fits to the simulated time series when using >6 seconds. GEDb provided overall the highest fits, while SSD was most sensitive to the amount of time. These differences can be attributed to differences in implementation of the temporal filters applied prior to eigendecomposition.

The final parameter test was the sensitivity to the number of electrodes. The simulated dipole time series were projected to 256 electrodes (layout taken from the standard EGI net), and random subsets of electrodes were used. The procedure was repeated 40 times per number of electrodes (each time selecting a different random subset). This simulation was run using four signal strengths. Results in Figure 4e show that the time course reconstruction accuracy increased to 50-70 electrodes. On the other hand, the simulated signal amplitude was a stronger determinant of R^2^ than was the number of electrodes. This demonstrates that having more electrodes is useful, but having a strong generating source is much more important. Though not a surprising conclusion, it highlights that simply having more data does not guarantee better results.

### Task-related EEG data

In the final set of analyses, an empirical task-related dataset was used, one subject randomly selected from a published dataset (Cohen, 2015). A single subject is included as proof-of-principle illustration in empirical data. The spatial filters were computed based on data from 0-700 ms relative to stimulus onset (at 512 Hz, this corresponds to 358 time points), when task-related dynamics can be expected. Results are presented in Figure 5.

**Figure 5.**
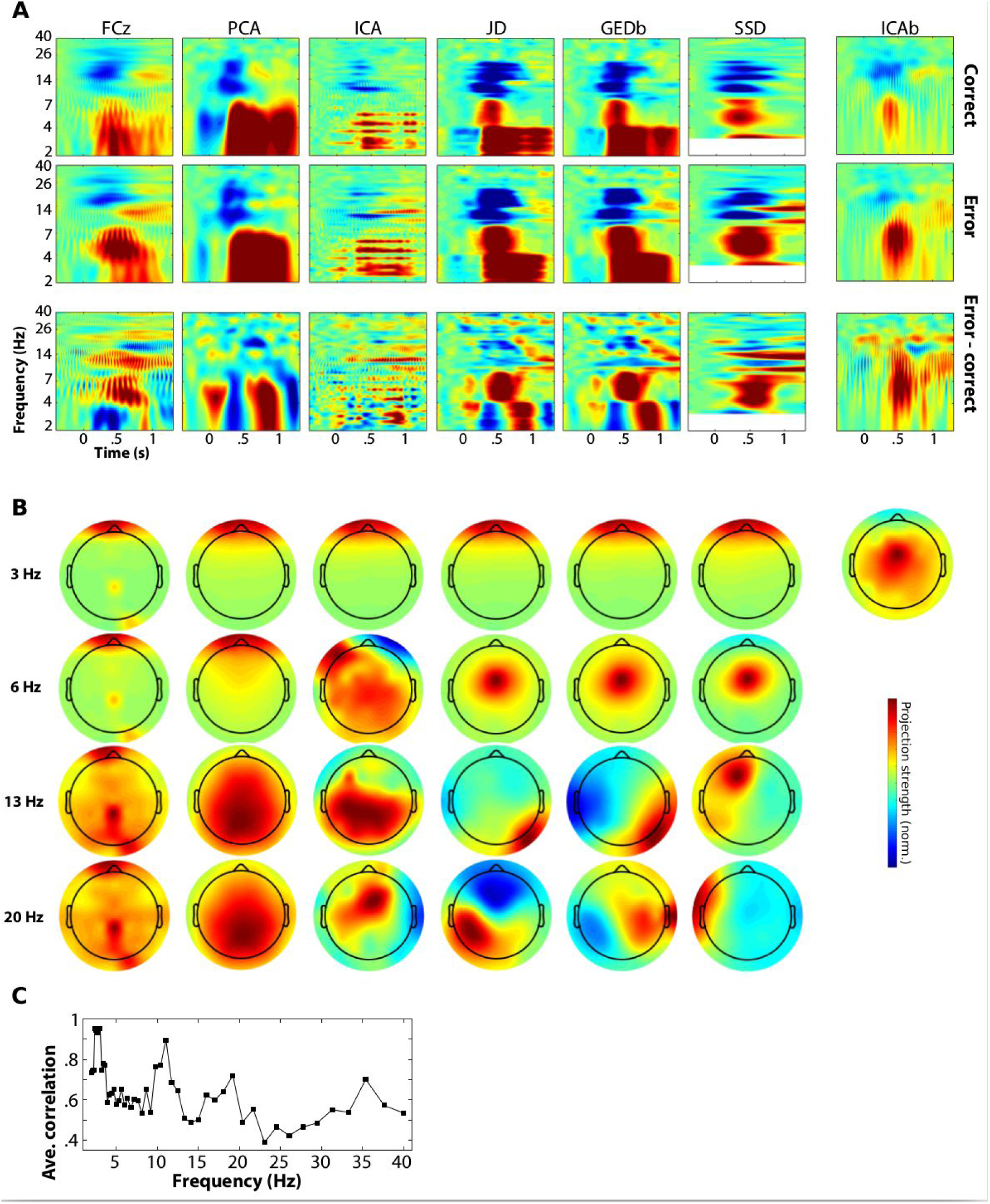
Results from task-related data (N=1). Spatial filters were applied at each frequency, and the power time course was extracted (top row shows correct trials, middle row shows error trials, bottom row shows their difference). Panel A shows the time-frequency power plots for each spatial filter and for electrode FCz. The rightmost column shows results from computing ICA from the broadband data and selecting a component with a midfrontal topography. Panel B shows topographical maps for selected frequencies, and panel C shows the average inter-map correlations across frequencies.

There are several noteworthy aspects of the results. First, the temporal filter kernel was too narrow for the single-electrode data—as evidenced by the ringing artifacts in the time-frequency plots—but was sufficient for the spatially filtered components. Second, the topographies differed not only across frequencies, but also across spatial filters. This indicates that different topographical regions contributed to the same time-frequency plot, and that different spatial filters were sensitive to different features of cortical network dynamics. One example is the stronger increase in delta-band power in several spatial filters compared to data from FCz. Inspection of the topographical maps revealed that this low-frequency power increase was from anterior electrodes, and likely reflects eye-movement artifacts (other than manual trial rejection, the EEG data were not cleaned using, e.g., ICA). It is therefore also interesting to note that the spatial filters at most other frequencies suppressed the oculomotor artifacts, despite having non-zero power at higher frequencies (evidenced by their representation in the electrode power plots).

Third, ICA following narrow bandpass filtering produced poor results. A single ICA decomposition based on the broadband data was then performed (“ICAb” in Figure 5), with a component selected for analysis based on having a midfrontal topography. Time-frequency results were more interpretable, although there were some filter ringing artifacts.

### Example application of spatial filtering

In this section, an application of spatial filtering for dimensionality reduction and SNR enhancement is illustrated, using the empirical data shown in Figure 5. It has previously been reported that midfrontal theta and posterior upper-alpha (~10-15 Hz) interact following response errors, and that these large-scale network fluctuations predict next-trial behavioral performance (Cohen and van Gaal, 2013).

Based on this a priori expectation, two components were extracted from the data using GEDb, one to isolate midfrontal theta (center frequency of 6 Hz) and one to isolate posterior alpha (center frequency of 13 Hz) (see Figure 6a for topographical projections). Spatial filters were computed using data from all 250 trials (using data from all trials improves the filter quality and eliminates biases from condition comparisons). Time-frequency power dynamics from these components were then computed, separately for correct and error trials, and can be seen in Figure 6b. Time-domain Granger prediction was then applied to the broadband time-domain signals, and the results indicate enhanced directed connectivity from the theta component to the alpha component following errors (Figure 6c). The same Granger prediction analysis applied to the electrodes with maximal topographical energy (see starred electrodes in Figure 6a) showed a somewhat different pattern of findings, with a later peak. The “true” effect in empirical data is unknown, but the increased accuracy of GEDb over single-electrode shown in Figures 2 and 3 suggests that the components-based connectivity result is a better indicator of the underlying neural dynamics. Firm conclusions about neurocognitive dynamics should not be made based on a single dataset; nonetheless, the important point is the illustration of how spatial filtering for component isolation and data reduction can be used in practice.

**Figure 6.**
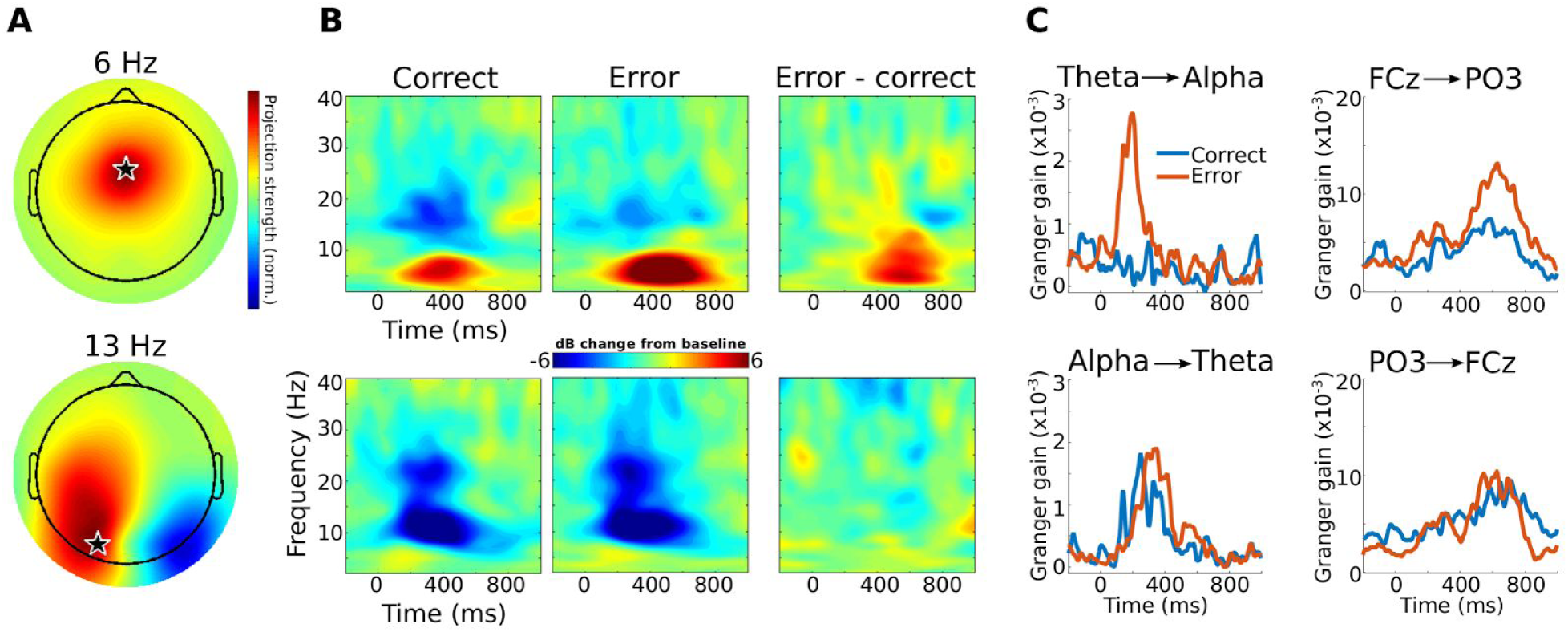
Example illustration of how spatial filtering for dimensionality reduction might be used in practice. Two components were extracted for theta and alpha. Topographical maps of the filter projections (panel A) and time-frequency power plots (panel B) for each component are illustrated. Time-domain Granger prediction (panel C) was applied to investigate how these components interact. Stronger theta**→**alpha directed connectivity after errors was observed, consistent with enhanced top-down signaling. In this panel, “theta” and “alpha” refer to the broadband data generated by applying spatial filters optimized for theta and alpha activity; the time series data were not bandpass filtered. The righthand column of panel C shows the same analysis applied to data from electrodes FCz and PO3, which were selected because they were the spatial maxima of the components (see starred locations in panel A).

## Discussion

### Spatial filtering in cognitive electrophysiology

The dominant uses of spatial filtering in cognitive electrophysiology are to attenuate sources of noise (e.g., using ICA during preprocessing), estimate anatomical localization (e.g., dipole-fitting or beamforming), facilitate functional connectivity analyses (e.g., Laplacian or source-space analyses), and, as highlighted here, as a dimensionality-reduction technique. This paper highlighted a few methods that, when thoughtfully constructed, can enhance SNR, isolate components in multichannel data, and alleviate the need to perform possibly suboptimal electrode selection.

Spatial filtering techniques such as the ones reviewed here are not widely used in the cognitive electrophysiology literature, although they are widely used in other disciplines, for example in the brain-computer-interface literature. There are several advantages of incorporating spatial filtering in cognitive electrophysiology research, the first two of which were demonstrated in this paper. First, spatial filters can be used to increase SNR. Here, the filters were optimized to maximize power at a given frequency band, but spatial filters can also be designed for other goals, such as maximizing a correlation with a behavioral or experiment variable (Dähne et al., 2014; Dmochowski et al., 2015). The improvement in SNR is clearly demonstrated in this study by comparing SNR values between the various spatial filters and the best-electrode “filter” in Figures 2-4.

Second, spatial filters can be used as a dimensionality-reduction tool, by allowing researchers to analyze a small number of components rather than a large number of electrodes. This was illustrated here by comparison with the best-electrode approach, in which an algorithm that selected the electrode with the largest power made suboptimal selections when the data contained noise. Electrode selection can be difficult and involve uncertainty and subjectivity. Components extracted from spatial filters might also facilitate cross-subjects comparisons by accounting for individual differences in anatomical and functional anatomy. For example, different individuals might have peak alpha activity at different electrodes (due to differences in cortical folding, electrode placement, etc.), but defining and then analyzing an alpha component for each subject does not require electrode-averaging or individual electrode-picking.

Third, spatial filters can be used to identify patterns in EEG that might be difficult to observe in unfiltered data. This is a primary motivation for applying spatial filters in machine learning applications, such as “decoding” features of sensory stimuli (King et al., 2014). Identifying spatial patterns in EEG might also be useful for low-SNR components, such as the visual C1 component, which can be difficult to identify in individual subjects based on electrode-selection (Proverbio et al., 2007).

Finally, spatial filtering can be used as a multivariate technique to identify large-scale networks defined by condition- or time-period differences (de Cheveigné and Parra, 2014).

#### Which method to use?

As is typical in neuroscience data analysis methods, no single method is superior in all contexts. Different methods are designed with different goals in mind, and there is little sense in assuming that any particular method should be optimal for situations outside the original intended goal. That said, a few generalities can be made based on the results described above.

JD had high sensitivity to recover the simulated time series, but in some cases this may lead to overfitting noise. Furthermore, the bias filter as implemented here (de Cheveigné and Parra, 2014) was phase-dependent and assumes sinusoidal stationarity. Both of these qualities seem suboptimal for isolating neural oscillations, which are often non-stationary and non-sinusoidal (Jones, 2016). However, the bias vector need not be a pure sine wave; it can take any number of forms, such as time-varying stimulus properties, data from an EMG electrode, or any other continuous variable.

GEDb performed best in most of the evaluations implemented here. GEDb does not suffer from phase- or filter kernel-dependencies because it is based on the covariance of the observed data from a narrow frequency range. Because there are no assumptions about the characteristics of the signal other than the temporal frequency used in narrowband filtering, GEDb is simple yet robust, which is a good characteristic of data analysis methods.

SSD had high SNR in many simulations. The accuracy of reconstructing the simulated time series, however, was lower than from GEDb or JD. Furthermore, high accuracy required longer data epochs (Figure 4d). This difference is attributable to the temporal filters applied in the SSD code that were less adept at capturing the temporal nonstationarities, which are contained in the spectral side-lobes.

PCA fared relatively poorly, in some cases even worse than the best-electrode approach (e.g., Figure 2A1). This should not be surprising: The use of PCA for isolating a single component for analysis or for removing artifacts has been empirically observed to under-perform in several other publications (Delorme et al., 2012; Haufe et al., 2014; Jung et al., 2000). There are two theoretical explanations for this. First, PCA returns eigenvectors that are pairwise orthogonal, which is arguably too strong and nonphysiological a constraint for multichannel brain recordings. Second, PCA has no minimization objective that is comparable to GED—PCA will maximize variance along particular directions, but the maximum directions of variance do not necessarily correspond to the patterns of variance that are scientifically relevant (de Cheveigné and Parra, 2014).

However, one should keep in mind that the poor performance of PCA refers to the use of PCA to isolate a single component; there are other applications of PCA that are valid and useful in neural data. For example, PCA can be used to reduce data dimensionality, e.g., from 128 channels to 60 components to facilitate ICA decomposition. In such cases, the goal is to define a still-high dimensional subspace of the data that can subsequently be decomposed into nonorthogonal components.

ICA also performed comparatively poorly. One key difference between ICA and the GED-based filters considered here is the objective of the filter: The goal of ICA is to decompose a multichannel signal into a set of independent components, while the goal of GED is to identify components that maximize a variance-power ratio between two features of the data (the two covariance matrices). Thus, GED is a “guided” source separation method whereas ICA is a blind source separation method. It is sensible that strong a priori knowledge about a component will lead to more accurate identification of that component.

Many techniques based on GED of two covariance matrices differ only in minor ways. Therefore, rather than advising to use one specific method or one specific set of parameters, it would behoove interested researchers to understand the general principle of using eigendecomposition to design linear spatial filters (Tomé, 2006) and then custom-tailor appropriate techniques for specific applications. For this reason, “blackbox” blind source separation techniques should be avoided in cognitive electrophysiology data. The temporal and spectral non-stationarities, in combination with noise, lead to a large risk of uninterpretable results (see Figure 5). Some level of human expertise and visual inspection is crucial to obtain suitable and interpretable results.

There are many more spatial filters and variants of filters than were evaluated here. Anatomical localization methods, for example, were not considered here, in part because distributed localization techniques can be thought of as dimensionality expansion methods rather than dimensionality reduction methods. Relatedly, the surface Laplacian, though beneficial in many circumstances (Kayser and Tenke, 2015), does not reduce the dimensionality of the data, which was the goal of the spatial filters applied here. In practice, a fruitful approach is first to consider the goal of spatial filtering, and then to find and optimize a particular spatial filter.

Relatedly, there are also dimensionality-reduction techniques that are not based on GED (Cunningham and Ghahramani, 2015). More mathematically sophisticated methods that incorporate iterative or nonlinear algorithms may produce more accurate results in some cases. However, simple linear methods have several advantages: they tend to be easy and fast to implement; they tend to be robust to nonstationarities, noise, and parameter settings; and they require less expertise to use appropriately. Nonetheless, the omission of iterative or nonlinear spatial filters in this paper is not a rejection of their usefulness; instead, the point was to highlight that spatial filters are powerful, insightful, and easy to incorporate into data analysis protocols.

#### How to select filter parameters?

Analysis parameter selection requires careful consideration, as illustrated in Figure 4. It should be kept in mind that the purpose of that figure was not to define a set of correct and incorrect parameters, but instead to highlight the sensitivity of filters to their parameters. Optimal parameters in one situation may be suboptimal in a different situation. For example, if SSD is used to identify alpha activity and if the “noise” covariance matrix is defined from data bandpass filtered in the theta band, the spatial filter may reflect suppression of theta activity rather than enhancement of alpha activity; more likely, it will contain a mixture of both, stymying an easy interpretation.

To the extent that features of the data can be known a priori and simulated, it is advisable to determine appropriate parameters based on simulations. Alternatively, parameters can be optimized in a subset of data and then applied to the rest of the data.

### Interpretation, practical use, and conclusions

The term “source” is widely used in electrophysiology with widely different interpretations. It sometimes refers to a physical location, sometimes to a latent component, and sometimes to cognitive process. In the case of spatial filters like the ones evaluated here, it is best to think of “source” as statistically isolated components of the data. A single component may be driven by an anatomically restricted neural generator, or it may reflect the coordinated activity of spatially distributed networks. A priori expectations and inspection of topographical maps can be used as evidence consistent with localized or distributed generators, but it is important to keep in mind that anatomical information is not considered when designing any of the filters used here.

For task-related data, and for comparison of activity over different frequencies, one must interpret the results carefully. Activity of different frequencies in a single time-frequency plot might come from very different topographical—and thus, neuroanatomical—regions, as illustrated in Figures 3 and 5. Whether this is problematic depends on the specific application. One can imagine generating a result like Figure 5 to determine the important spatial-temporal-spectral features of the data, and then perform follow-up analyses at frequencies/electrodes empirically demonstrated to be task-relevant. Perhaps a better approach is illustrated in Figure 6, in which a priori hypotheses were used to define two components based on expected spectral characteristics; the time series of those two components were then analyzed instead of time series from 64 electrodes. For this approach, it is advantageous to define the spatial filter based on bandpass filtered data and then apply the spatial filter to the broadband data (Cheveigné et al., 2015)as recommended by (Cheveigné et al., 2015); this facilitates subsequent analyses such as time-frequency power and Granger prediction (which should not be applied to bandpass filtered data).

Electrophysiological data contain many dimensions of information that vary in complex and dynamical ways over time, frequency, and space, and as a function of cognitive processes. Furthermore, one is often faced with limited and noisy data. Completely blind source separation methods with no expert human intervention or guidance are unlikely to produce interpretable results. However, strong a priori expectations about the relevant spectral-temporal-spatial characteristics of the data can be used in combination with spatial filtering and data reduction techniques to increase SNR and facilitate data analysis. This may become increasingly important as the study of electrical activity underlying human brain function becomes increasingly sophisticated, and as the number of electrodes used steady increases.

